# CTRR-ncRNA: A Translation-oriented Knowledgebase for Cancer Resistance and Recurrence Associated Non-coding RNAs

**DOI:** 10.1101/2022.08.01.502291

**Authors:** Tong Tang, Xingyun Liu, Rongrong Wu, Li Shen, Shumin Ren, Bairong Shen

## Abstract

Cancer therapy resistance and recurrence (CTRR) are the dominant causes of death in cancer patients. Recent studies have indicated that non-coding RNAs (ncRNAs) can not only reverse the resistance to cancer therapy but also are crucial biomarkers for the evaluation and prediction of CTRR. Herein, we developed CTRR-ncRNA, a knowledgebase of CTRR-associated ncRNAs, aiming to provide an accurate and comprehensive resource for research involving association between CTRR and ncRNAs. Compared to most of the existing cancer databases, CTRR-ncRNA is focused on the clinical characterization of cancers, including cancer subtypes, as well as survival outcomes and response to personalized therapy of cancer patients. Information pertaining to biomarker ncRNAs has also been documented for the development of personalized CTRR prediction. A user-friendly interface and several functional modules have been incorporated into the database. Based on the preliminary analysis of genotype–phenotype relationships, universal ncRNAs have been found to be potential biomarkers for CTRR. The CTRR-ncRNA is a translation-oriented knowledgebase and it provides a valuable resource for mechanistic investigations and explainable artificial intelligence-based modelling. CTRR-ncRNA is freely available to the public at http://ctrr.bioinf.org.cn/.

## Introduction

Cancer patients may develop resistance to previously effective treatments, including chemotherapy, radiotherapy, and immunotherapy [1–3]. Investigations of the mechanisms underlying cancer therapy resistance and recurrence (CTRR) are made complicated by the presence of confounding factors, such as heterogeneous genetic backgrounds, as well as diversity in cancer cells and tumour microenvironments [1–4]. Reducing or eliminating the CTRR has been a tough challenge.

Protein-coding sequences only account for approximately 1.5% of the human genome, and the majority of the genome is associated with ncRNAs, such as microRNAs (miRNAs), long ncRNAs (lncRNAs), and circular RNAs (circRNAs) [5, 6]. The evidence accumulated so far indicates that ncRNAs play crucial roles in the modulation of treatment resistance [7, 8]. For instance, miR-144-3p can regulate the resistance of lung cancer to cisplatin by targeting the gene encoding nuclear factor erythroid 2–related factor 2 (Nrf2) [9]. By inhibiting the wnt2/β-catenin signaling pathway, LINC00968 can attenuate drug resistance in breast cancer [10]. Knockdown of hsa_circ_0081143 has been shown to reverse the cisplatin resistance by targeting the miR-646/CDK6 pathway in gastric cancer [11]. In addition, more and more evidence shows that ncRNAs have significant functions in regulating radiotherapeutic and immunotherapeutic resistance. Radiotherapeutic resistance in breast cancer can be modulated by miR-139-5p [12]. lncRNA CCAT2 and miR-191 promote resistance to radiotherapy in human oesophageal carcinoma cells and prostate cancer, respectively [13, 14]. The circRNA circFGFR1 can promote anti-PD-1 resistance by sponging miR-381-3p in non-small cell lung cancer [15]. As indicated in multiple reports, ncRNAs have been regarded as one of the most promising molecules to reverse therapeutic resistance [16, 17].

During the cancer treatment, some cancer cells may escape from their original locations and continue to survive. These cells may subsequently grow into large enough tumours to cause cancer recurrence, thereby increasing the complexity and difficulty of cancer treatment [18]. Previous research supports that ncRNAs could prove to be biomarkers for diagnosis, treatment, and prognosis for recurrence in different types of cancer. For example, miRNA-125b could be a prognostic marker in recurrent hepatocellular carcinoma as its levels are significantly lower in the early stages than in the late stages of recurrence [19]. LncRNA CASC2a is identified as not only a prognosis biomarker but also a potential therapeutic target in the early recurrence of bladder cancer [20]. These data confirm the significance of ncRNAs in cancer recurrence [21–24].

A large number of biological experiments and clinical trials have proven that many ncRNAs are involved in reversing treatment resistance and prevention of cancer recurrence [13, 25–27]. However, a public resource for ncRNAs related to CTRR is not unavailable. Indeed, some previously created databases focus on drug resistance, but they ignore the information regarding radiotherapeutic and immunotherapeutic resistance [28–30]. The data documented in these databases are derived from basic biological experiments without clinical records. With the accumulation of evidence for the correlation between ncRNAs and clinical traits observed in cancer patients, it is becoming increasingly necessary to build a database that integrates knowledge from biological and clinical domains. Such a database would be valuable for systematic modelling and personalized treatment of CTRR [31].

In recent decades, a variety of techniques are applied to identify biomarkers for CTRR. Most of the biomarker discovery methods are based on wet-lab experiments, such as small RNA sequencing and RT-qPCR assay [32, 33]. Computational approaches are also developed that are complementary to or integrated with experimental methods for efficient biomarker identification [34]. For example, network smoothed T-statistics-based biomarker detection [35] and network vulnerability-based identification of potential biomarkers [36, 37]. A competitive endogenous RNA (ceRNA) network has also been built to discover the critical position through network topology analysis for biomarker discovery [38]. However, until now, very few computational methods or knowledgebases have been developed that are specific for CTRR ncRNA biomarkers discovery.

To this end, we developed the CTRR-ncRNA knowledgebase to provide resources for CTRR associated ncRNA biomarker discovery and customized analysis of the association between CTRR and ncRNAs. In this database, we have emphasized on the regulatory mechanism of ncRNAs, the clinical characterization of ncRNAs associated with CTRR, and annotation of ncRNA biomarkers.

## Data collection and processing

### Data collection and knowledge extraction

We started by searching for literature in the PubMed database using the search terms listed in File S1. As a result, 3998 publications describing an association between CTRR and non-coding RNAs were obtained. These publications were further filtered using the following exclusion criteria, 1) Research involving specimens other than human cell lines or tissues, 2) Resistance alteration caused by epigenetic modification of ncRNAs, 3) Resistance altered by the interaction between drugs and ncRNAs, and 4) Reviews, meta-analyses, and bioinformatics analyses without experimental validation. Post filtering, biological and clinical information regarding ncRNAs and CTRR was manually curated.

For cancer therapy resistance, we retrieved the symbols of ncRNAs, cancer names, therapeutic methods, upstream regulatory factors and downstream targets of ncRNAs, biomarkers and their clinical applications such as diagnosis, treatment, prognosis, the impact of ncRNA status, such as wildtype, knockdown, or overexpression on the treatment resistance. In addition, we paid much attention to the collection of clinical information related to cancer therapy resistance, including the difference in expression of an ncRNA between patients and healthy people, clinical sample size, sex ratio, and age ranges. Finally, the experimental details regarding the use of cell lines vs. patient’s tissues, the description of ncRNA’s role in cancer therapy resistance were collected. The following information was collated for the annotation of cancer recurrence 1) Cancer recurrence type, 2) Overall survival of patients with altered ncRNAs expression, 3) Ratios of recurrence; 4) Sex ratios of recurrence, 5) Age ratios of recurrence, and 6) The differential expression between the recurrence and control groups.

#### Standardization of ncRNA nomenclature

To reduce the confusion caused by ncRNA synonyms, the names of miRNA and circular RNA were standardized based on the names used in the miRbase [48] and circBase [49], respectively. miRNA nomenclature has changed dramatically in recent years. The miRbase v22 (https://www.mirbase.org/) was utilized as the reference for the standardization of miRNA nomenclature. For example, hsa-miR-24 used in the previous studies [50, 51] was renamed as “hsa-miR-24-3p” in CTRR-ncRNA. If the miRNAs were not included in miRbase v22, their original names were kept unchanged, such as “hsa-miR-128” [52]. Similar rules were applied for circular RNA nomenclature, e.g., human circEIF6 [53] was standardized as hsa_circ_0060060 in accordance with the circBase. Both human circRNAs Cdr1as [54] and CDR1as [55] were standardized as hsa_circ_0001946. Currently, there are no unified nomenclature rules for lncRNA, hence, we have used lncRNA names from the original resources. In addition, we have documented the body site of the cancers in the database as per International Classification of Diseases 11th to provide information for the personalized cancer diagnosis, prognosis, and treatment.

### Online database implementation

The CTRR-ncRNA knowledgebase is available online at ctrr.bioinf.org.cn. The web framework is Flask 1.0, the database is MySQL 5.7, the user interface framework is Bootstrap 4.4, and the website is deployed using Ngnix. It has been tested in Microsoft Edge, Mozilla, Chrome, and Safari browsers. A user-friendly interface is provided.

### Cancer-ncRNA network construction and characterization

The cancer-ncRNA network was constructed based on the specific association between different ncRNAs and cancer types. The different cancers were considered as phenotypes and the CTRR related ncRNA-status was regarded as the genotypes. Their relationships were analysed at the systems biology level. The ncRNAs were classified as specific or universal, based on their relationship with cancers, i.e., if the association between the ncRNA and the cancer is unique, the ncRNA was defined as specific, otherwise, it was classified as universal. Cytoscape was used to visualize the Cancer-ncRNA network [56]. The scale-free model [57] was applied to evaluate and characterize the Cancer-ncRNA network.

## Data content and usage

### CTRR-ncRNA database statistics

In Total, 367 ncRNAs associated with cancer therapy resistance in 79 cancer types and 46 ncRNAs associated with cancer recurrence in 24 different cancer types are included in the CTRR-ncRNA knowledgebase. All the experimentally validated CTRR associated ncRNAs, by qRT-PCR, western blot, a luciferase reporter assay, etc., are included in the database. The ncRNA entries related to resistance in the database include 497 miRNAs (65%), 248 lncRNAs (32%), and 24 circRNAs (3%). Additionally the ncRNA entries related to recurrence in the database include 33 miRNAs(69%) and 13 lncRNAs (31%). These statistics can be found on the website at http://ctrr.bioinf.org.cn/. The overview of the database construction and implementation is illustrated in Figure 1.

**Figure 1.**
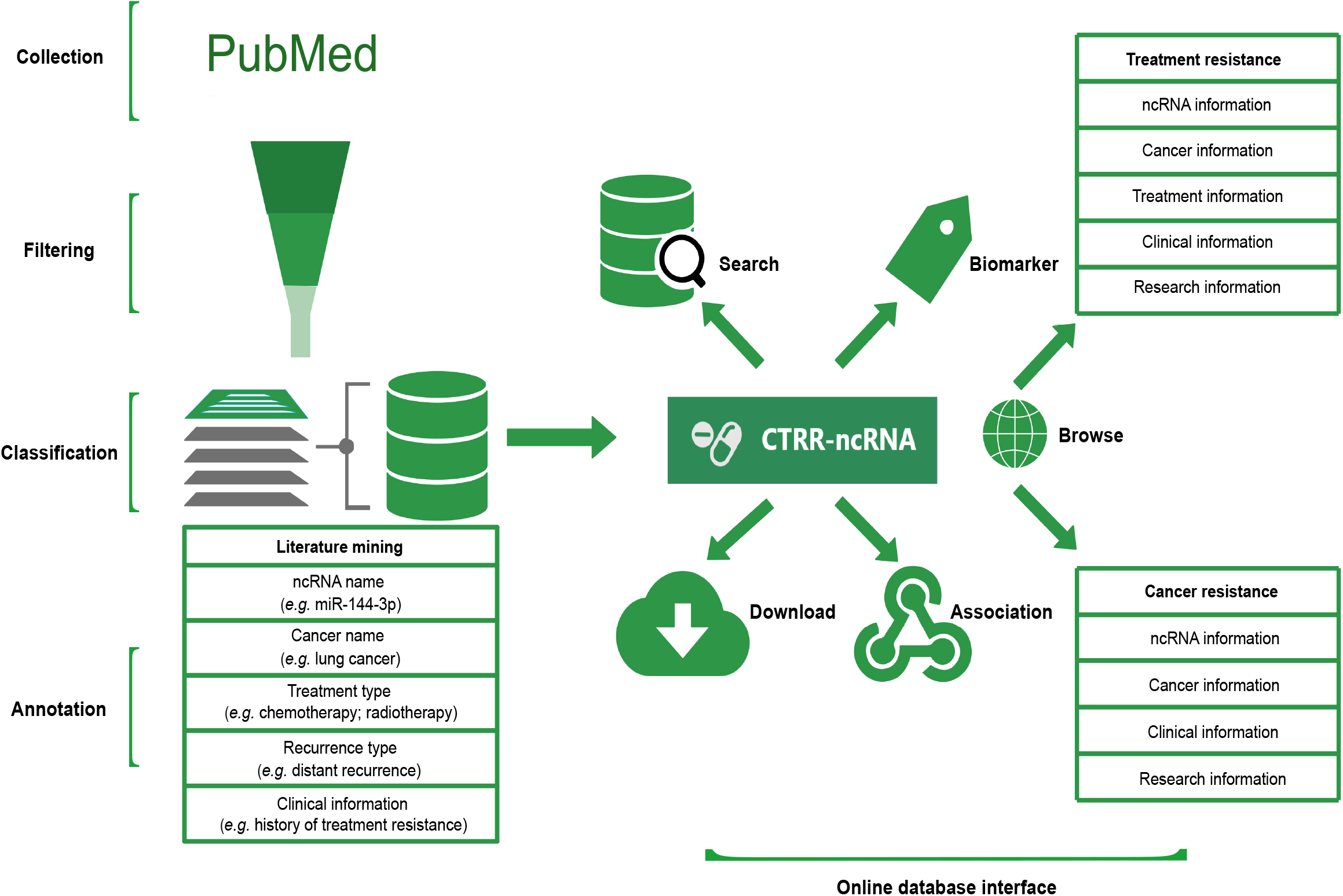
An overview of data collection, filtering, classification, annotation, and online database interface.

### Web pages and functions

The CTRR-ncRNA knowledgebase includes multiple web pages including the home page (Figure 2A and 2B) which provide access to information on cancer therapy resistance and cancer recurrence, database search, data download, and contact information.

**Figure 2.**
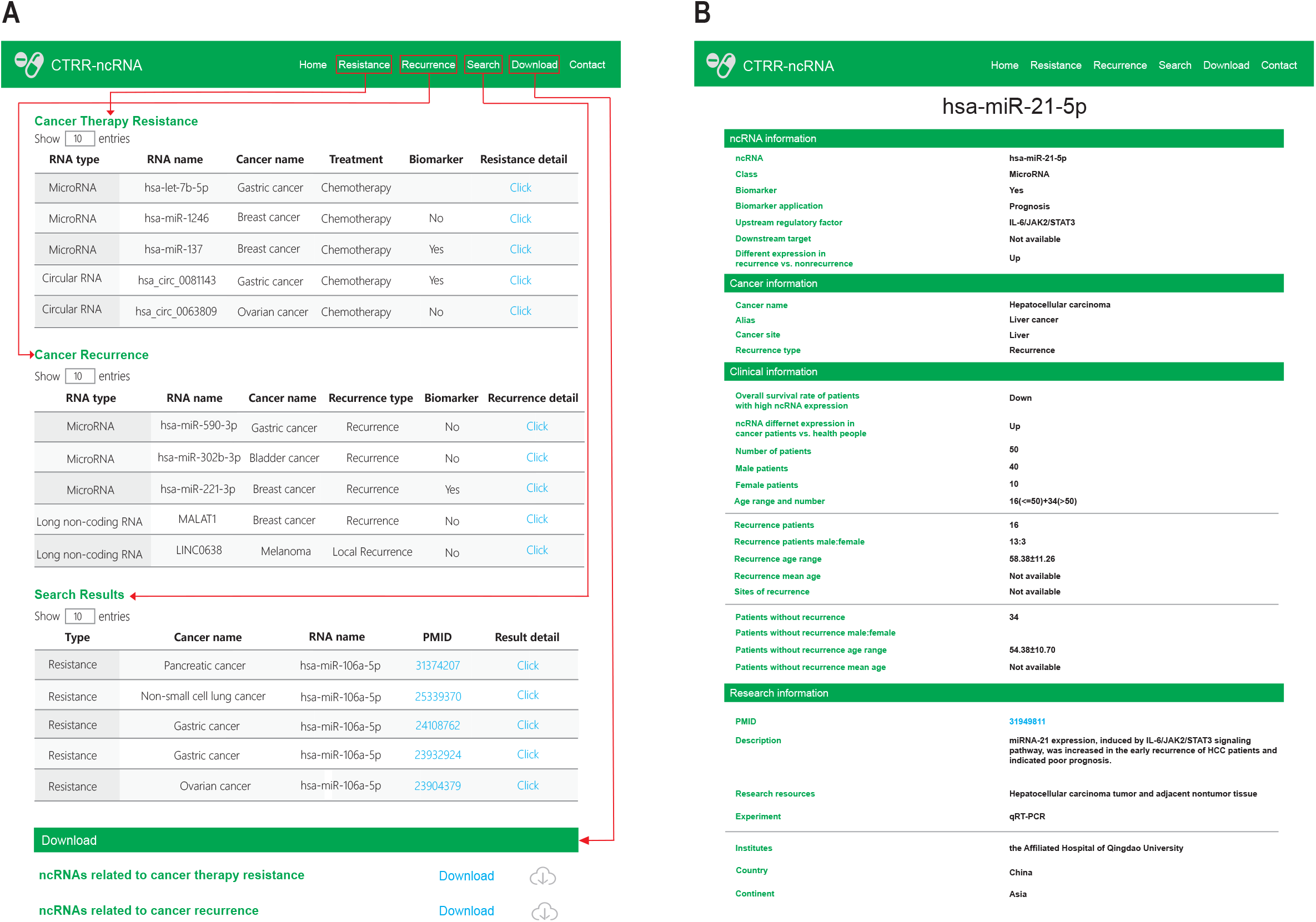
A schematic design of CTRR-ncRNA. A. The ‘Resistance’, ‘Recurrence’, ‘Search’ and ‘Download’ modules allow the users to browse, search or download ncRNAs. B. Details of an ncRNA entry.

### The home page

The schematic diagram of the association between ncRNAs and CTRR is illustrated at the top of the page. A brief introduction of the CTRR-ncRNA database and its features is described at the bottom of the page.

### The page for cancer therapeutic resistance

This page is designed to access information regarding cancer therapy resistance and the associated ncRNAs. Users can extract the information by ncRNAs, cancers, biomarkers, and clinical applications, personalized treatments, upstream regulatory factors and downstream targets of ncRNAs, CTRR associated ncRNA status. The entry details can be displayed by clicking on the “Click” button to check the information about the ncRNAs, cancer types, treatment, clinical information, and details about the original research.

### The page for cancer recurrence

We have provided detailed clinical information regarding cancer recurrence cases including overall survival of patients with abnormal ncRNAs expression and the clinical information of cancer-recurrence patients. Cancer patients’ overall survival values are correlated with the expression levels of the associated ncRNAs. Therefore, the CTRR-ncRNA would prove to be helpful for the personalized prognosis of cancer recurrence cases.

### The page for searching

On this page, a quick fuzzy search function is provided for the users for efficient data retrieval. The contents of the CTRR-ncRNA knowledgebase have been classified structurally and users can quickly screen the data of interest by ncRNA names, cancer names, treatment or recurrence types, and biomarker applications.

### Download and contact pages

The Download page is available for the users to download all entries of the CTRR-ncRNA knowledgebase. A tutorial for the usage of the database is provided on the Contact page.

### Comparison with the existing databases

To date, several databases are available that have collected the ncRNAs related to drug resistance. They are well-structured and include information regarding ncRNAs and drug targets, but no databases were established about ncRNAs associated with CTRR. CTRR-ncRNA collected the 26 ncRNA records regarding radiotherapeutic and immunotherapeutic resistance which are useful for the research of different treatment. The advantages of our database as compared to other related databases are presented in Table 1.

Personalized diagnosis, treatment, and prognosis of cancer is the key issue in this era of precision medicine. It is necessary to combine the personalized clinical information with molecular omics profiles. However, most of the existing ncRNA databases are focused on the relationship between ncRNAs and drugs, but lack the information associated with clinical attributes this limits the application of these databases for clinical use. For example, ncRNA Drug Targets Database (NRDTD) has primarily collected descriptive information related to diseases, drugs, and ncRNAs, without clinical information or clinical applications [28]. It also lacks the description of the relationship between diseases, drugs, and ncRNA. Compared to the NRDTD database, the NoncoRNA database has a more detailed description of the relationship between diseases, drugs, and ncRNA [30]. However, it does not include details about the association between ncRNAs and therapy resistance or their clinical applications.

The CTRR-ncRNA contains information about the relationship between ncRNA expression and CTRR. The upstream regulatory factors and downstream targets of ncRNAs, the clinical information of patients including sample sizes, sex ratios, age ranges, and expression of ncRNAs are included in the present database. We have also paid special attention to the ncRNA biomarkers, which have application in the CTRR related diagnosis, treatment, and prognosis.

In the design and implementation of our database, we have focused on the integration of clinical information with the classification of CTRR. The cancer recurrence was classified as local, distant, local and distant, early, and metastatic recurrence. The clinical information regarding overall survival and the ncRNAs expression was extracted from the literature.

### Cancer-ncRNA network characterization

To investigate the genotype-phenotype relationship in CRTT, we first constructed the cancer-ncRNA association network for cancer therapy resistance, as shown in Figure 3. The hsa-miR-150-5p and lncRNA MALAT1 are universal ncRNAs, which are associated with many types of cancers. The hsa-miR-150-5p is reported as a prognostic biomarker for recurrent ovarian cancer, and lncRNA MALAT1 is an indicator for recurrence of gallbladder cancer [39]. As presented in Figure S1, most of the universal ncRNAs are reported as biomarkers in cancer therapy resistance, such as hsa-miR-31-5p [40–42], HOTAIR [43], MALAT1 [44], and LINC00152 [45]. The scale-free model (*P*(*K*)~ *K^-λ^*) that fitted to our cancer-ncRNA network is presented in Figure 4. The *λ* is 2.04 and 2.20 for the cancer-miRNA network and cancer-lncRNA network respectively. The *λ* value between two and three indicates that the ncRNA nodes connected to many cancer types are crucial [46], and many experimental results support that universal ncRNAs are more likely to be biomarkers in CTRR [47].

**Figure 3.**
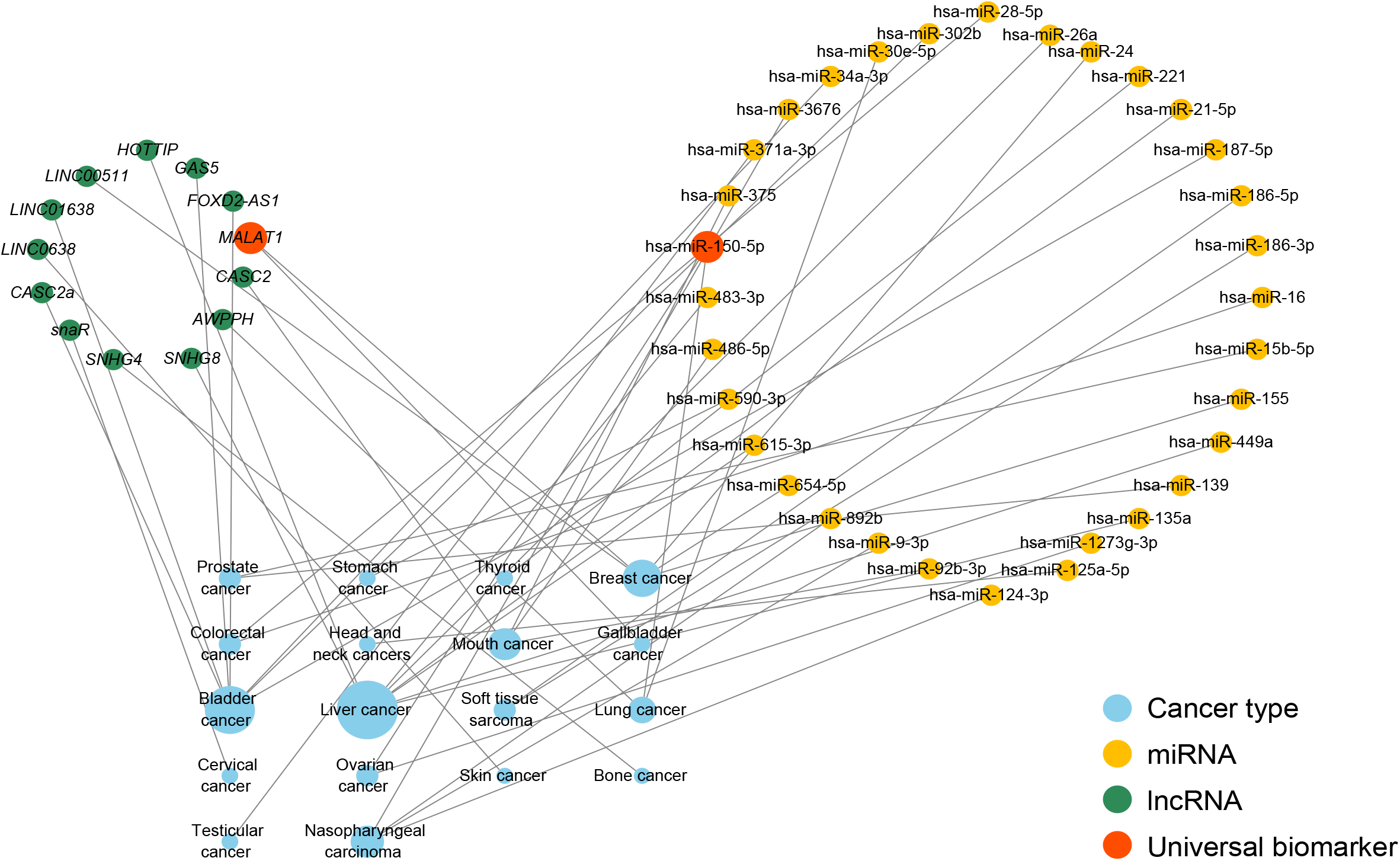
An overview of the cancer–ncRNA network associated with cancer recurrence. The yellow circles represent miRNAs, the green circles represent lncRNAs, and the orange circles represent universal ncRNAs. The blue circles represent cancer types and the size of the blue circles is proportional to the number of cancer-associated ncRNAs.

**Figure 4.**
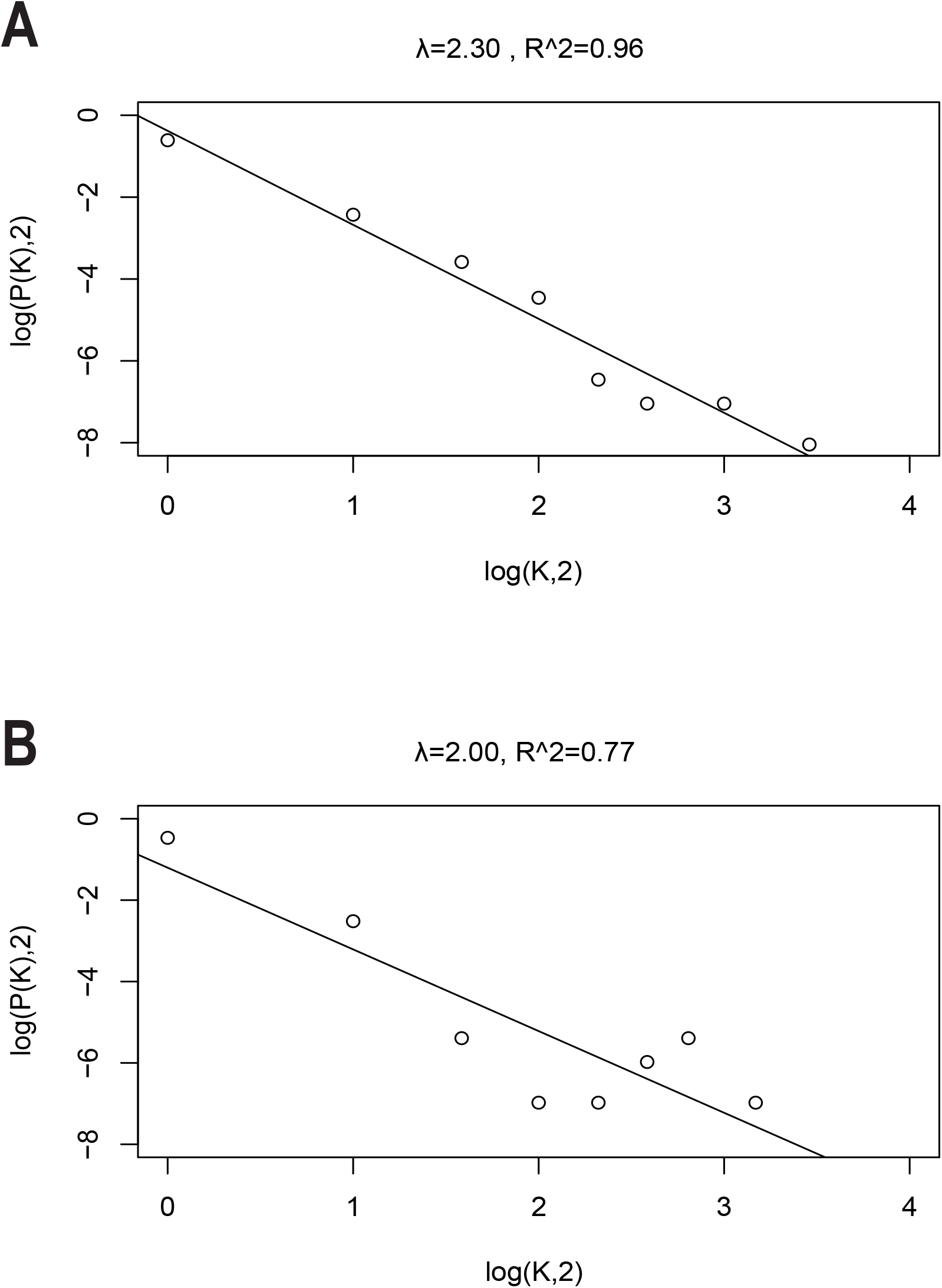
Correlations and distribution of miRNAs and lncRNAs related to the database. A. Scale-free feature of miRNAs related to cancer therapy resistance. B. Scale-free feature of lncRNAs related to cancer therapy resistance.

## Conclusion

The CTRR-ncRNA can provide information for mapping the personalized relationship between ncRNA, cancer, and clinical information, for the pattern discovery in the genotype-phenotype relationship, and for supporting the clinical decision.

## Data availability

CTRR-ncRNA is publicly available at http://ctrr.bioinf.org.cn/.

## CRediT author statement

**Tong Tang**: Data curation, Formal analysis, Methodology, Visualization, Writing - original draft. **Xingyun Liu**: Data curation, Methodology. **Rongrong Wu**: Data curation, Formal analysis. Li Shen: Visualization, Writing - review & editing. **Shumin Ren**: Formal Analysis, Writing - review & editing. **Bairong Shen**: Conceptualization, Methodology, Supervision, Project administration, Writing - review & editing. All the authors have read and approved the final manuscript.

## Acknowledgments

We thank the members of the Institutes for Systems Genetics for the discussion and assistance with the data collection. This work was supported by the National Natural Science Foundation of China (Grant No. 32070671) and the regional innovation cooperation between Sichuan and Guangxi Provinces (Grant No. 2020YFQ0019).

## Supplementary material

**File S1 Search terms used for screening PubMed database**

**Figure S1 An overview of the cancer–ncRNA network in cancer therapeutic resistance**

