## Supplementary figures and images for "CTRR-ncRNA: A Translation-oriented Knowledgebase for Cancer Resistance and Recurrence Associated Non-coding RNAs"

### Figure S1

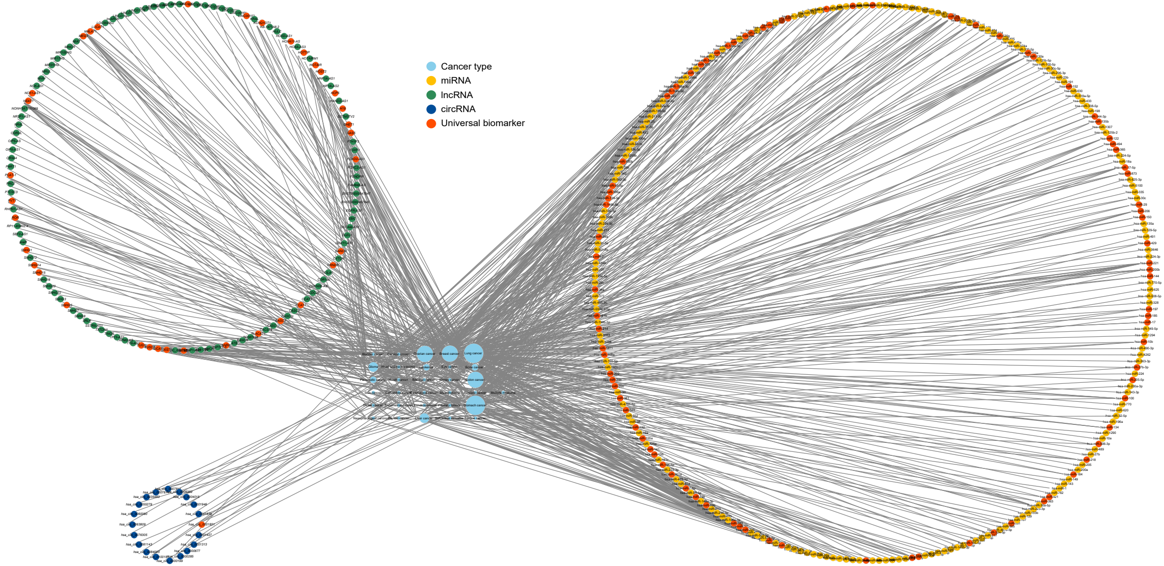
