## Supplementary material for "CTRR-ncRNA: A Translation-oriented Knowledgebase for Cancer Resistance and Recurrence Associated Non-coding RNAs": File S1

(resistan*[ti] OR relaps*[ti] OR recurren*[ti]) AND (cancer*[tiab] OR tumor*[tiab] OR carcinoma*[tiab] OR neoplasm*[tiab] OR glioma*[tiab] OR meningiomas*[tiab] OR retinoblastoma*[tiab] OR hepatoblastoma*[tiab] OR Cholangiocarcinoma*[tiab] OR cholangiocarcinoma*[tiab] OR neuroblastoma*[tiab] OR endometrial*[tiab] OR lymphoma*[tiab] OR sarcoma*[tiab] OR osteosarcoma*[tiab] OR leukemia*[tiab] OR histiocytoma*[tiab] OR rhabdomyosarcoma*[tiab] OR myeloma*[tiab] OR melanoma*[tiab] OR pheochromocytoma*[tiab]) AND (non-coding RNA*[tiab] OR ncRNA*[tiab] OR transfer RNA*[tiab] OR tRNA*[tiab] OR ribosomal RNA*[tiab] OR rRNA*[tiab] OR miRNA*[tiab] OR microRNA*[tiab] OR siRNA*[tiab] OR piRNA*[tiab] OR snRNA*[tiab] OR snoRNA*[tiab] OR Extracellular RNA*[tiab] OR exRNA*[tiab] OR scaRNA*[tiab] OR circular RNA*[tiab] OR circRNA*[tiab] OR lncRNA*[tiab] OR long non-coding RNA*[tiab] OR lincRNA*[tiab]) AND (“1990/01/01”[DP] : “2020/1/1”[DP])
